# BiGPICC: a graph-based approach to identifying carcinogenic gene combinations from mutation data

**DOI:** 10.1101/2023.02.06.527327

**Authors:** Vladyslav Oles, Sajal Dash, Ramu Anandakrishnan

## Abstract

Genome data from cancer patients represents relationships between the presence of a gene mutation and cancer occurrence in a patient. Different types of cancer in human are thought to be caused by combinations of two to nine gene mutations. Identifying these combinations through traditional exhaustive search requires the amount of computation that scales exponentially with the combination size and in most cases is intractable even for cutting-edge supercomputers.

We propose a parameter-free heuristic approach that leverages the intrinsic topology of gene-patient mutations to identify carcinogenic combinations. The biological relevance of the identified combinations is measured by using them to predict the presence of tumor in previously unseen samples. The resulting classifiers for 16 cancer types perform on par with exhaustive search results, and score the average of 80.1% sensitivity and 91.6% specificity for the best choice of hit range per cancer type. Our approach is able to find higher-hit carcinogenic combinations targeting which would take years of computations using exhaustive search.

## 1 Introduction

Multi-hit theory of carcinogenesis states that it takes combinations of gene mutations to initiate carcinogenesis in humans^1^. Clinical studies and a body of mathematical models established that the size of such combinations (hits) can range from 2 to 9 depending on the cancer types^2–6^. However, most computational efforts to find carcinogenic mutations focus on finding individual “driver mutations” responsible for carcinogenesis^7–10^. While these mostly mutational frequency- and signature-based methods identify driver mutations associated with increased risk of cancer, these mutations by themselves cannot cause cancer.

Cancer-screening methods based on genetic predisposition rely heavily on identifying driver mutations. However, some people with a genetic predisposition may never get cancer, while others get cancer, having accumulated more mutations with age. For example, women under the age of 20 with a BRCA1 mutation are very unlikely to get breast cancer; moreover, 28% of all women with BRCA1 mutation never get the cancer^11^. Similar statistics can be observed for Li Fraumeni syndrome in men^12–15^. These observations provide strong evidence that cancer in a patient is caused not by an individual driver gene mutation but rather by a combination of gradually accumulated gene mutations. Though there are several other factors responsible for cancer such as epigenetic modifications, tumor environment, and adaptive evolution, carcinogenesis is primarily a result of genetic mutations.

Different combinations of gene mutations can cause cancer of the same type but with different etiologies and pathologies representing different subtypes. To design and develop individualized and precision drugs for treating an individual with cancer, we need to identify the carcinogenic combinations of gene mutations in that patient. Since current computational approaches mostly focus on singular driver gene mutations, they fall short in the context of precision drug discovery. The human genome 𝒢 is comprised of 𝒢 ≈ 20, 000 genes^16^, with each gene potentially hosting dozens of mutations. Even if the task of identifying carcinogenic combinations of *mutations* is replaced with a simpler task of identifying carcinogenic combinations of *genes* (i.e. combinations of genes that can harbor jointly carcinogenic mutations), solving it for the combinations of size *h* > 4 would be infeasible due to the need of enumerating all 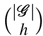 possibilities (e.g. over 0.26 · 10^20^ combinations even for *h* = 5). In this work, we propose an approach that can efficiently find carcinogenic gene combinations even for large values of *h*.

### 1.1 Exhaustive search methods

A recent body of work^17–19^ explored all possible gene combinations to identify those responsible for cancer based on tumor and normal samples from cancer patients. They mapped the task to a weighted set cover problem, where each combination is associated with a set of tumor samples in which it is jointly mutated and assigned weight based on its classification accuracy when used for differentiating between tumor and normal samples. They implemented a heuristic approximating the minimum weight set cover that requires 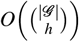 time for identifying a single combination of length *h*. This, however, is only feasible if *h* ≤ 4, as in the case of *h* > 4 their exhaustive search would take days to years on one of the fastest supercomputers in the world, Summit (see the calculations in Section 3.6).

Klein et al.^20^ developed another computational method for inferring combinations of gene mutations that cooperatively drive tumor formation in individual patients. Their parameter-heavy approach combines exhaustive search and heuristics when navigating the combinatorial space for the human genome, and can identify carcinogenic combinations of length *h* for *h* ≤ 6.

Methods like Dendrix^21^, Multi-Dendrix^22^, SuperDendrix^23^, CoMEt^24^ etc. employ exhaustive search after constraining the search space of human genome 𝒢 by breaking it up into gene sets with approximately mutually exclusive mutations. This only partially offsets the issue of exponential time complexity in *h*, and, to the best of our knowledge, such approaches has only been applied for *h* ≤ 5^23^ (see “Bioinformatics and Data processing” in the Supplemental therein). Additionally, requiring the mutual exclusivity in target mutations comes at a cost of potentially lower biological relevance, similarly to other parameter-based approaches to reducing the search space size.

### 1.2 Graph algorithms in cancer genomics

Graphs are combinatorial structures that model pairwise relationships between objects, e.g. protein-protein interactions (PPI) or mutations of genes in humans. In recent years, a number of studies have relied on graph-based analyses to identify combinations of cancer drivers^25–30^. A typical approach is to enrich a molecular network of choice using gene-sample mutation data, then analyze the resulting graph, usually by means of network propagation, to identify clusters of carcinogenic genes. For example, Leiserson et al.^26^ use a heat diffusion algorithm on PPI networks to find their mutated subnetworks, based on the gene mutation frequencies extracted from data. The identified communities (or modules) are hypothesized to represent carcinogenic gene combinations, and many of the corresponding proteins have a documented role in development of cancer. In a similar vein, Leung et al.^27^ employ a greedy search to identify co-mutated communities in gene networks. Cho et al.^25^ rely on gene neighborhoods in functional networks in combination with mutation information.

Unlike the above studies, Iranzo et al.^31^ directly use the graphical structure of mutation data to examine community structure in cancer drivers. The approach of this work is similar to ours in the idea of leveraging the topology of gene-sample mutations to learn about group interactions of carcinogenic genes, although their methodology is different. Another distinction is that Iranzo et al. study the ~1% of the human genome that has known associations with cancer, while our work aims to discover novel carcinogenic combinations by analyzing the full genome.

### 1.3 Our contribution

Here we present BiGPICC (Bipartite Graph Partitioning for Identifying Carcinogenic Combinations) — a parameter-free approach to finding multi-hit combinations based on the topology of mutation data. Analyzing mutation data is potentially more insightful for cancer genomics than using molecular networks, as the latter are simplified abstractions of gene or protein interactions. To the best of our knowledge, this is one of the first network-based algorithms working directly with gene-sample mutations.

We formulate the search in terms of a community detection problem on a bipartite graph representation of the data, and design and implement an algorithm for solving it. Our numerical experiments on Summit supercomputer for 16 cancer types demonstrate that it identifies combinations of comparable biological relevance to state-of-the-art results. At the same time, our approach is capable of efficiently identifying relevant (5+)-hit combinations, unavailable to the existing methods due to high computational cost. An additional advantage of out method is that it does not require any manual tuning due to the absence of parameters representing some form of domain knowledge, which means that it is readily available to work with a broad variety of datasets.

## 2 Methods

Our approach for identifying carcinogenic gene combinations based on the presence of their exonic, protein-altering mutations in tumor and normal tissues is outlined in Figure 1. Viewing the binary mutation data (same as used in^17–19^) as a bipartite graph, we iteratively partition it using community detection to find candidate gene combinations whose mutations tend to occur in tumor tissues, then filter out those with frequent mutations in normal samples. The final set of carcinogenic combinations is identified by selecting a minimum set cover of the tumor samples from the filtered candidate pool. To assess the relevance of the identified combinations, we use them to differentiate between tumor and normal samples on previously unseen data. A detailed description of these steps is provided below.

**Figure 1.**
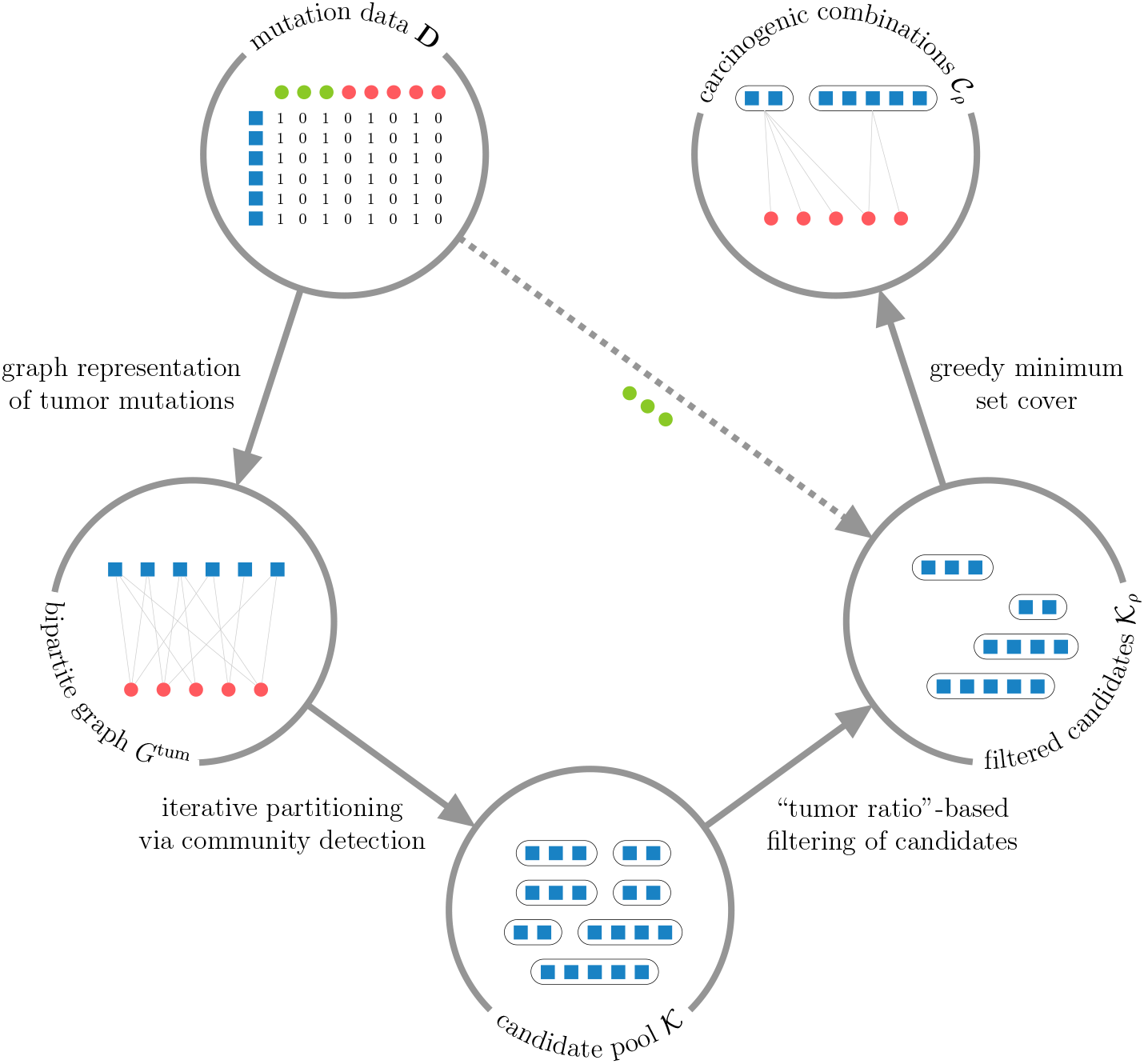
Outline of BiGPICC approach, where blue squares represent genes and green and red circles represent, respectively, normal and tumor samples. Starting from binary mutation data (**upper left**), we construct a bipartite graph of mutations in tumor samples (**left**), use its structure to find gene combinations frequently mutated in tumor samples (**bottom**), retain only the combinations rarely mutated in normal samples (**right**), and select their smallest non-redundant subset as the output combinations (**upper right**). The four processing stages of our approach, shown as solid arrows in counterclockwise order starting from the upper left, are described respectively in Sections 2.1 through 2.4.

### 2.1 Graph formulation using binary mutation data

Let 𝒢 and 𝒮 denote the sets of genes and samples in the data, respectively. The input data contains information about whether a mutation of gene *g* was observed in sample *s* for every gene-sample pair (*g, s*) ∈ 𝒢 × 𝒮. Specifically, the data is stored in a binary | 𝒢 | × | 𝒮 | matrix **D** whose entries are 1 if a mutation of the corresponding gene was observed in the corresponding sample, and 0 otherwise (note that it does not differentiate between mutations within the same gene). This information can be equivalently represented as an unweighted bipartite graph on the vertices of two distinct classes 𝒢 and 𝒮, where an edge connects some *g* ∈ 𝒢 and *s* ∈ 𝒮 if and only if the corresponding mutation has been observed. We denote such a graph *G*, and notice that the adjacency matrix of *G* is the symmetric block matrix 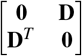. The presence of both normal and tumor samples in the data means that the vertex class 𝒮 is itself partitioned into two subclasses, denoted 𝒮 ^norm^ and 𝒮 ^tum^ respectively, with opposing significance to the problem. To isolate the information about gene mutations in tumor tissues, we consider the graph *G*^tum^ := *G*[𝒢 ⊔𝒮 ^tum^], the induced subgraph of *G* on the vertices 𝒢 ⊔𝒮 ^tum^.

Recall that a *community* is a subset of graph vertices that are more densely connected with one another than with the rest of the graph, according to some metric. Any community *C* in *G* (or *G*^tum^) consists of the gene component *C*_𝒢_ :=*C* ∩ 𝒢 — a combination of genes whose mutations tend to occur together, and the sample component *C*_𝒮_ :=*C ∩* 𝒮 (or *C ∩* 𝒮 ^tum^, respectively) — the samples in which these genes co-mutate the most. Because bipartite communities of multiple vertices must contain vertices from both classes to allow for internal connections, both *C*_𝒢_ and *C*_𝒮_ are non-empty whenever |*C*| > 1.

For some gene *g* ∈ 𝒢, let *M*(*g*) ⊂ 𝒮 denote the samples in which a mutation of *g* has been observed. Furthermore, for a gene combination *C*_𝒢_, denote the samples in which these genes are jointly mutated as *M*(*C*_𝒢_) := ∩∈*C*𝒢 *M*(*g*). Analogously, we define the tumor samples with a mutation of *g* as *M*^tum^(*g*):= *M*(*g*) ∩ 𝒮 ^tum^, and with a joint mutation of *C*_𝒢_ — as *M*^tum^(*C*_𝒢_) := *M*(*C*_𝒢_) ∩ 𝒮 ^tum^. If *C*_𝒢_ is carcinogenic, its joint mutation is thought to explain some number of tumor samples from the data, so *C*_𝒢_ must be fully connected to the non-empty set *M*^tum^(*C*_𝒢_) in the graph representation of data. For any unrelated gene *g*^*′*^ ∈ 𝒢 \ *C*_𝒢_, the connectivity between genes *C*_𝒢_ ∪ {*g*^*′*^} and samples *M*^tum^(*C*_𝒢_) is expected to be less dense, because a mutation of *g*^*′*^ is unlikely to appear in every sample from *M*^tum^(*C*_𝒢_) by chance unless it is mutated in most of 𝒮 ^tum^ (which would suggest insufficient. For example, if *C*_𝒢_ is jointly mutated in |*M*^tum^(*C*_𝒢_)| = 10 tumor samples and *g*^*′*^ is mutated in half of all the tumor samples (i.e. *M*^tum^(*g*^*′*^) = 0.5|𝒮 ^tum^|), the probability of its mutation to appear in all of *M*^tum^(*C*_𝒢_), i.e. of *g*^*′*^ forming a community with *C*_𝒢_ ⊔*M*^tum^(*C*_𝒢_), is

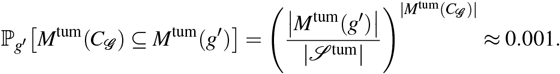

It follows that in the presence of a sufficient number of tumor samples any carcinogenic gene combination *C*_𝒢_ is expected to correspond to a community *C*_𝒢_ ⊔ *M*^tum^(*C*_𝒢_) in *G*^tum^. At the same time, the converse is not necessarily true — because carcinogenic combinations are assumed to jointly mutate in predominantely tumor samples, not every community in *G*^tum^ has its gene component as a carcinogenic combination. For example, if tumor samples constitute only a small fraction of the samples with a joint mutation of *C*_𝒢_ (i.e. |*M*^tum^(*C*_𝒢)_ |_≪_ | *M*(*C*_𝒢_) |), then *C*_𝒢_ is unlikely to be a carcinogenic combination even despite *C*_𝒢_ ⊔*M*^tum^(*C*_𝒢_) being a community in *G*^tum^. Therefore, the task of finding carcinogenic combinations of gene mutations can be cast as a problem of identifying communities with some desired degree of tumor prevalence in their sample components.

### 2.2 Community detection

We use the *Constant Potts Model* (CPM) to formally define the notion of community structure in a graph. CPM was proposed as an alternative to the commonly-chosen *modularity* approach to alleviate the issue of inconsistent communities across different scales^32^. For the case of *G*^tum^, an unweighted bipartite graph on the vertex classes 𝒢 and 𝒮 ^tum^, CPM formalizes its partition into disjoint communities 𝓅 (i.e.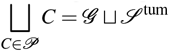) as desirable if it maximizes the partition quality

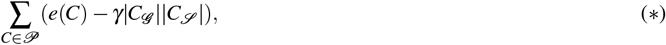

where *e*(*C*) is the total number of internal edges in community *C* (that is, between *C*_𝒢_ = *C ∩* 𝒢 and *C*_𝒮_ = *C ∩* 𝒮 ^tum^), and *γ* ∈ [0, 1] is the so-called resolution parameter. This parameter can be viewed as the density of internal connectivity required from a set of vertices *C* for it to qualify as a community — or, more specifically, for *C* to positively contribute to the partition quality (*). Notice that whenever gene component or sample component of *C* are empty, its contribution to (*) is zero.

To identify communities within a graph, we rely on the Leiden algorithm^33^, chosen for its guarantees on the community connectivity and speed. In particular, the Leiden algorithm converges to a partition in which all subsets of all communities are guaranteed to be locally optimally assigned. The Leiden algorithm starts by viewing every vertex as a separate community, and then alternates between moving nodes between communities, refining the partition, and aggregating communities into single vertices to reduce the graph size, until no further improvement to the chosen partition quality (in our case, (*)) can be made.

Because the possible number of carcinogenic drivers in a combination is known or hypothesized for many cancer types, we are interested in finding communities whose gene component size *h* := |*C*_𝒢_ | is within a certain range *l* ≤ *h* ≤ *u* as obtained from the literature. To enforce this size constraint, we iteratively refine the identified communities using the Leiden algorithm until the size of their gene component does not exceed *u*, the maximum possible number of carcinogenic drivers. Specifically, if a community *C* in *G*^tum^ has gene component *C*_𝒢_ that is too large, we refine it by first extending its sample component to be 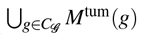 to include all relevant tumor samples, then applying the Leiden algorithm to the resulting subgraph 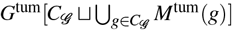 to partition *C*_𝒢_ based on their mutations. Every time the Leiden algorithm is run on a (sub)graph induced by genes *C*_𝒢_ and samples *C*_𝒮_, the CPM resolution parameter is set to the connectivity density of this (sub)graph, 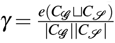, requiring the communities to be internally connected at least as densely as the (sub)graph itself. After filtering out the the communities whose gene component is too small from the results, the gene components of the remaining communities are considered candidates for carcinogenic combinations.

The above approach to partitioning *G*^tum^ to identify candidate gene combinations is described formally in Procedure 1. Because the Leiden algorithm is randomized, every call of Procedure 1 may yield a different set of candidate gene combinations. To increase the likelihood of carcinogenic combinations to appear among the results, we perform multiple iterative partitioning passes via Procedure 1 and combine obtained sets of candidate combinations *𝒦*^•^ into the joint candidate pool *𝒦*.

#### Procedure 1

Iterative Partitioning

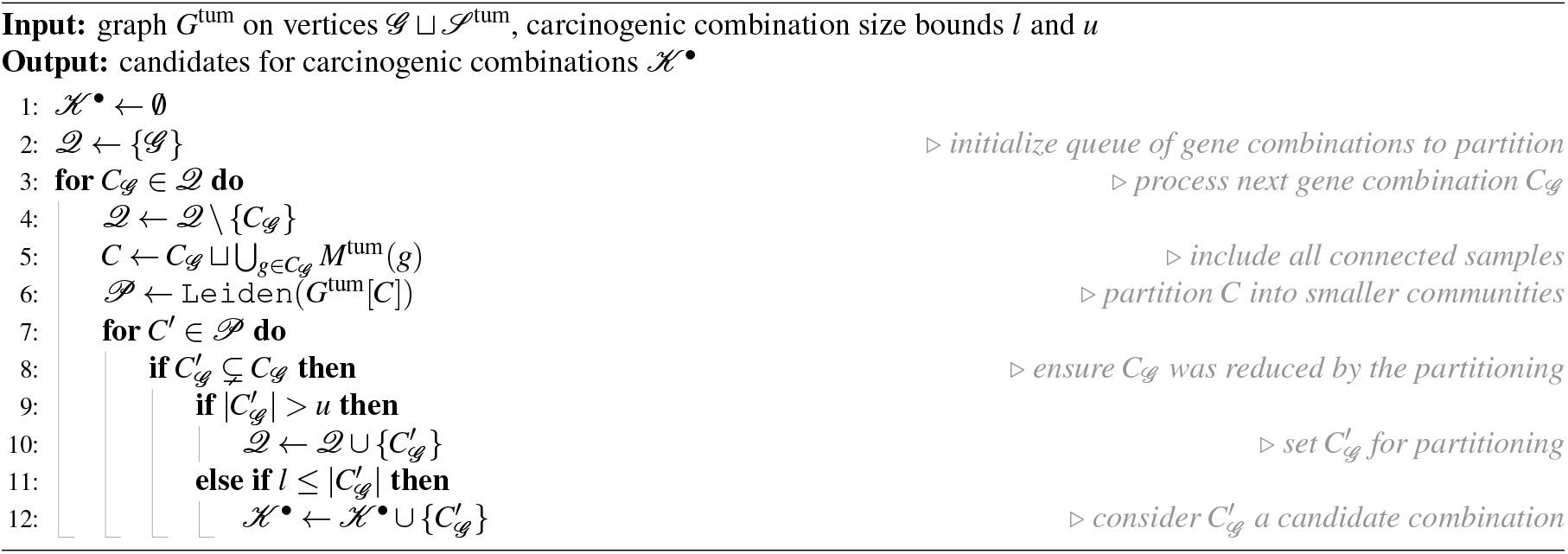

The iterative partitioning passes are independent of each other and can be performed in distributed fashion. We run them in parallel on the Summit supercomputer, relying on the implementation of Leiden algorithm from the Python library `leidenalg`.

### 2.3 Filtering of candidates

For a candidate combination *C*_𝒢_ ∈ *𝒦*, define its *tumor ratio* as the share of tumor samples among all those in which *C*_𝒢_ is jointly mutated, 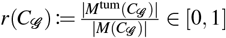 Naturally, true carcinogenic combinations are expected to have a higher tumor ratio as they can harbor mutations turning a normal sample into a tumor one. We impose a threshold *ρ* ∈ [0, 1] on the tumor ratio of candidate combinations, so that only “sufficiently carcinogenic” candidates *𝒦*_*ρ*_ := {*C*_𝒢_ ∈ *𝒦* : *r*(*C*_𝒢_) ≥ *ρ}* (notice that *𝒦*_0_ = *𝒦*), i.e. those with tumor ratio at least *ρ*, are considered when selecting the final set of carcinogenic combinations from the pool. For example, the choice of *ρ* = 1 implies considering only candidate combinations with no joint mutations in normal samples. This particular choice however is likely to result in implausible carcinogenic combinations due to the possibility of non-mutagenic drivers behind carcinogenesis and potential inaccuracies of genomic data.

### 2.4 Minimum set cover

For a reasonable choice of *ρ*, the set of tumor samples explained by the “sufficiently carcinogenic” candidates, 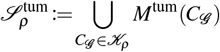 is expected to coincide with 𝒮^tum^ assuming that every tumor there is caused by a joint mutation of some gene combination from 𝒢 whose size is in the hypothesized range.

Given the threshold *ρ*, we select carcinogenic combinations as a subset of *𝒦*_*ρ*_ whose joint mutations explain 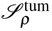 most concisely, by constructing a *minimum set cover* of 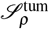 using sets of tumor samples from {*M*^tum^(*C*_𝒢_) : *C*_𝒢_ ∈ *𝒦*_*ρ*_. Specifically, we employ a greedy heuristic that iteratively chooses a candidate combination from *𝒦*_*ρ*_ to cover the majority of yet unexplained samples in 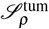 on each step (see Procedure 2). The size of the resulting set of carcinogenic combinations *𝒞*_*ρ*_ is guaranteed to approximate the size of the true solution within the factor of ln *m*, where *m* :=max {|*M*^tum^(*C*_𝒢_) | : *C*_𝒢_ ∈ *𝒦*_*ρ*_ is the biggest number of tumor samples explained by a “sufficiently carcinogenic” candidate^34^.

#### Procedure 2

Greedy Minimum Cover

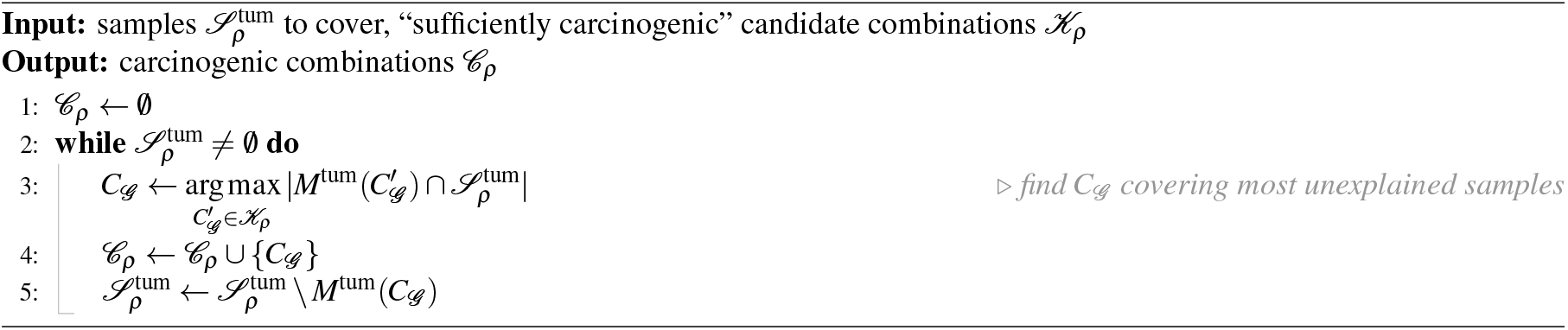

### 2.5 Algorithmic complexity

The time complexity of Procedure 1 is defined by the number and complexity of the calls to the Leiden algorithm it makes. Let *n*_0_, *n*_1_, … denote the numbers of genes in the inputs to these calls in chronological order. In particular, *n*_0_ = |𝒢 | as the first Leiden algorithm call takes the whole graph *G*^tum^ as the input. For any *k* > 0, the input genes of the *k*-th call are a strict subset of the input genes of some previous call and therefore *n*_*k*_ ≤ *n* _*j*_ − 1 for some *j < k*. Also, if every *i < k* satisfies *n*_*i*_ ≤ |𝒢 |−*i* then *i* = *k* also does, as *n*_*k*_ ≤ *n*_*j*_ − 1 ≤ |𝒢 | − *j* − 1 ≤ |𝒢 | − *k*. Because the condition is satisfied for *k* = 1, we obtain by induction that *n*_*i*_ ≤ |𝒢 |−*i* for every *i*. It follows that the number of Leiden algorithm calls made by Procedure 1 is *O*(|𝒢 |), and because the *i*-th call takes *O*(|𝒢 |−*i*) genes and *O*(|𝒮 |) samples, its time complexity is 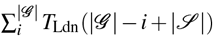, where *T*_Ldn_(*n*) is the runtime of the Leiden algorithm on a graph with *n* vertices.

Because partitioning of 𝒢 into combinations of size ≥ *l* produces at most 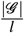 candidate combinations, their total number after calling Procedure 1 *p* times is bounded by | *𝒦* | = *O*(*p* 𝒢 *l*^−1^). Procedure 2 enumerates the remaining candidate combinations on each step, and terminates after at most 𝒮 ^tum^ steps. If a step covers only one new tumor sample, updating the count of yet unexplained samples covered by some candidate combination takes *O*(1) time. Viewing each step covering several new tumor samples as multiple single-sample steps then bounds the time complexity of Procedure 2 by *O*(|𝒮 ^tum^||*𝒦* |) = *O*(*p*|𝒮 ^tum^||𝒢 |*l*^−1^).

The resulting time complexity of BiGPICC pipeline is 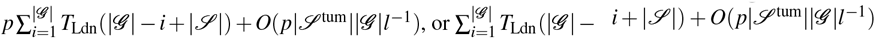 if the *p* calls to Procedure 1 are done in parallel. While it was proved that a CPM-optimal partition of an arbitrary *n*-vertex graph is reachable in *O*(*n*) steps of the Leiden algorithm, no upper bound on the number of steps, and therefore on *T*_Ldn_(*n*), is known. However, the algorithm was empirically shown to run in near-*O*(*n*) time on a variety of real-world and generated graphs of up to 10^7^ vertices^33^. If assuming linear runtime of the Leiden algorithm, the BiGPICC time complexity becomes 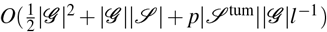.

The mutation data is represented as a |𝒢 |×|𝒮 | adjacency matrix. Candidate combinations identified in the community detection step are a superset of all BiGPICC outputs and span the total of *O*(*p*|𝒢 |) non-unique vertices. Therefore, the method’s memory complexity is *O*(|𝒢 ||𝒮 | + *p*|𝒢 |).

### 2.6 Learning threshold for the tumor ratio

To choose the best value of *ρ*, we frame the BiGPICC pipeline as a classification problem by using the selected carcinogenic combinations *𝒞*_*ρ*_ to predict whether previously unseen samples are normal or tumor. Namely, a sample is classified as tumor if it has a joint mutation of any *C*_𝒢_ ∈ *𝒞*_*ρ*_, and as normal otherwise. Notice that an increase in the value of *ρ* improves the precision score of the classifier while also growing the size of *𝒞*_*ρ*_ (as the choice of *C*_𝒢_ ∈ *𝒦*_*ρ*_ at each step of Procedure 2 is narrowed). Because the number of gene combinations in *𝒞*_*ρ*_ corresponds to classifier’s complexity and therefore its propensity to overfit, 1 − *ρ* can be viewed as the amount of regularization in the training, a trade-off between the “carcenogenicity” (in the sense of high tumor ratio) and the generalizability of learned combinations.

Before BiGPICC can access the data, we remove 25% of the samples to serve as a test dataset 𝒮_test_ for the final model. On the remaining 75% samples 𝒮*\* 𝒮_test_, we employ 4-fold cross-validation to find the optimal value of hyperparameter *ρ*. Specifically, 𝒮 *\* 𝒮_test_ is partitioned into equal parts 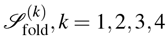 and in 4 separate scenarios the pipeline is run using the samples 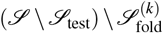 to produce *𝒞*_*ρ*_ for each value of *ρ* ranging from 0 to 1 with the increment of 0.01. The values of *ρ* are then assessed by averaging the performance of *𝒞*_*ρ*_ -based binary classifier on 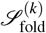 across the 4 scenarios. We use the *Matthews correlation coefficient* (MCC, also called the *phi coefficient*) — which takes values between -1 and 1 and is considered more informative than the commonly chosen F1 and accuracy scores^35^ — as a single performance metric. Because constructing *𝒞*_*ρ*_ and evaluating its performance for individual values of *ρ* can be done in parallel, the search for best *ρ* does not introduce a significant computational overhead. The value of *ρ* delivering highest mean MCC is then used to train the final classifier (by re-running BiGPICC pipeline) on the samples 𝒮 *\* 𝒮_test_ and report its performance on 𝒮_test_.

## 3 Results

### 3.1 Experimental setup

Our somatic mutation data was collected in mutation annotation format (MAF) from the original 2018 version of The Cancer Genome Atlas (TCGA) dataset for 16 cancer types using Mutect2 software. We identify a set of 331 matched blood-derived normal samples from all cancer types. We use the Variant Effect Predictor (VEP) to determine the location (intron, exon, UTR) and effect of these variants (synonymous/non-synonymous, missense/nonsense), and only consider protein-altering variants: non-synonymous, nonsense, and insertion/deletions in exons. For additional details about the protocols of data collection, see ^17^.

We apply BiGPICC to 16 datasets for different cancer types in 33 numerical experiments, each exploring a combination size *h* or its range for a cancer type according to existing literature. Each of the 16 datasets contains the same set of 331 normal tissue samples. Table 1 shows the hypothesized ranges of *h* and the dataset parameters for each cancer type. For each cancer type and multi-hit range, we run the community detection step (Procedure 1) *p* = 10,000 times to increase the probability of including all carcinogenic combinations to the candidate pool.

**Table 1.** Hit ranges required for carcinogenesis and data parameters for the 16 cancer types.

| Abbreviation | Cancer type | $h$, # hits | $\vert \mathcal{G}\vert $, # genes | $\vert \mathcal{S}\vert $, # samples | # mutations |
| --- | --- | --- | --- | --- | --- |
| BLCA | Bladder urothelial carcinoma | 4, 6-7, 7 | 18,511 | 699 | 519,225 |
| BRCA | Breast Invasive arcinoma | 3, 2-3, 6-7 | 19,411 | 1,242 | 837,207 |
| CESC | Cervical squamous cell carcinoma and endocervical adenocarcinoma | 5, 6-7 | 18,813 | 605 | 459,351 |
| COAD | Colon adenocarcinoma | 3, 4, 5, 6 | 19,487 | 716 | 665,932 |
| GBM | Glioblastoma multiforme | 2, 3 | 18,920 | 662 | 852,723 |
| HNSC | Head and neck squamous cell carcinoma | 4, 5 | 18,707 | 801 | 635,410 |
| KIRP | Kidney renal papillary cell carcinoma | 8 | 18,117 | 559 | 429,005 |
| LGG | Brain lower grade glioma | 3, 6 | 17,770 | 810 | 495,887 |
| LIHC | Liver hepatocellular carcinoma | 8 | 17,592 | 643 | 438,879 |
| LUAD | Lung adenocarcinoma | 3, 6-7 | 18,783 | 740 | 629,351 |
| LUSC | Lung squamous cell carcinoma | 5 | 18,454 | 636 | 488,441 |
| OV | Ovarian serous cystadenocarcinoma | 2, 6-7 | 18,814 | 658 | 671,888 |
| PRAD | Prostate adenocarcinoma | 6-7, 7 | 17,274 | 752 | 476,115 |
| SARC | Sarcoma | 5, 8 | 17,687 | 550 | 379,824 |
| THCA | Thyroid carcinoma | 5, 6 | 18,264 | 752 | 524,992 |
| UCEC | Uterine corpus endometrial carcinoma | 2, 6-7 | 19,889 | 826 | 911,903 |

### 3.2 Classification performance

In each numerical experiment we build a classifier based on the identified gene combinations of desired size to differentiate between tumor and normal samples. We use the MCC, specificity, sensitivity, and F1 metrics to highlight various aspects of our classifiers. Each of the numerical experiments performs the 4-fold cross-validation runs and chooses the best value of *ρ* based on the highest mean MCC across the 4 folds. We demonstrate the effect of the value of *ρ* on the MCC in individual numerical experiments in Figure 2. After using the chosen value of *ρ* to train the final classifier on 𝒢 ⊔ (𝒮 *\* 𝒮_valid_), we report its combination count and performance metrics on the previously unseen test samples 𝒮_test_ in Table 2. The spread of classification performance within the same cancer type is likely attributed to the biological relevancy of the chosen hit range in individual experiments. The variation in performance across different cancer types can be additionally explained by the number of available samples (e.g. whether it is sufficient to accurately distinguish between driver and passenger mutations) and their representativity of oncogenenic mutations as opposed to other factors (e.g. epigenetic or microenvironmental). In particular, the fact that the 4-hit BLCA classifier in ^19^ (see Figure 9 therein) performs as poorly as ours but a better sensitivity is achieved in ^20^ (Table 5) and ^18^ (Figure 4) using different hit ranges indicates that gene combinations whose mutations are driving BLCA are not predominantly 4-hit. Analogously, a significantly better performance of PRAD classifiers in ^17–19^ suggests that the hit range of 6 ≤ *h* ≤ 7 chosen for PRAD in our study does not cover all relevant drivers.

**Table 2.**
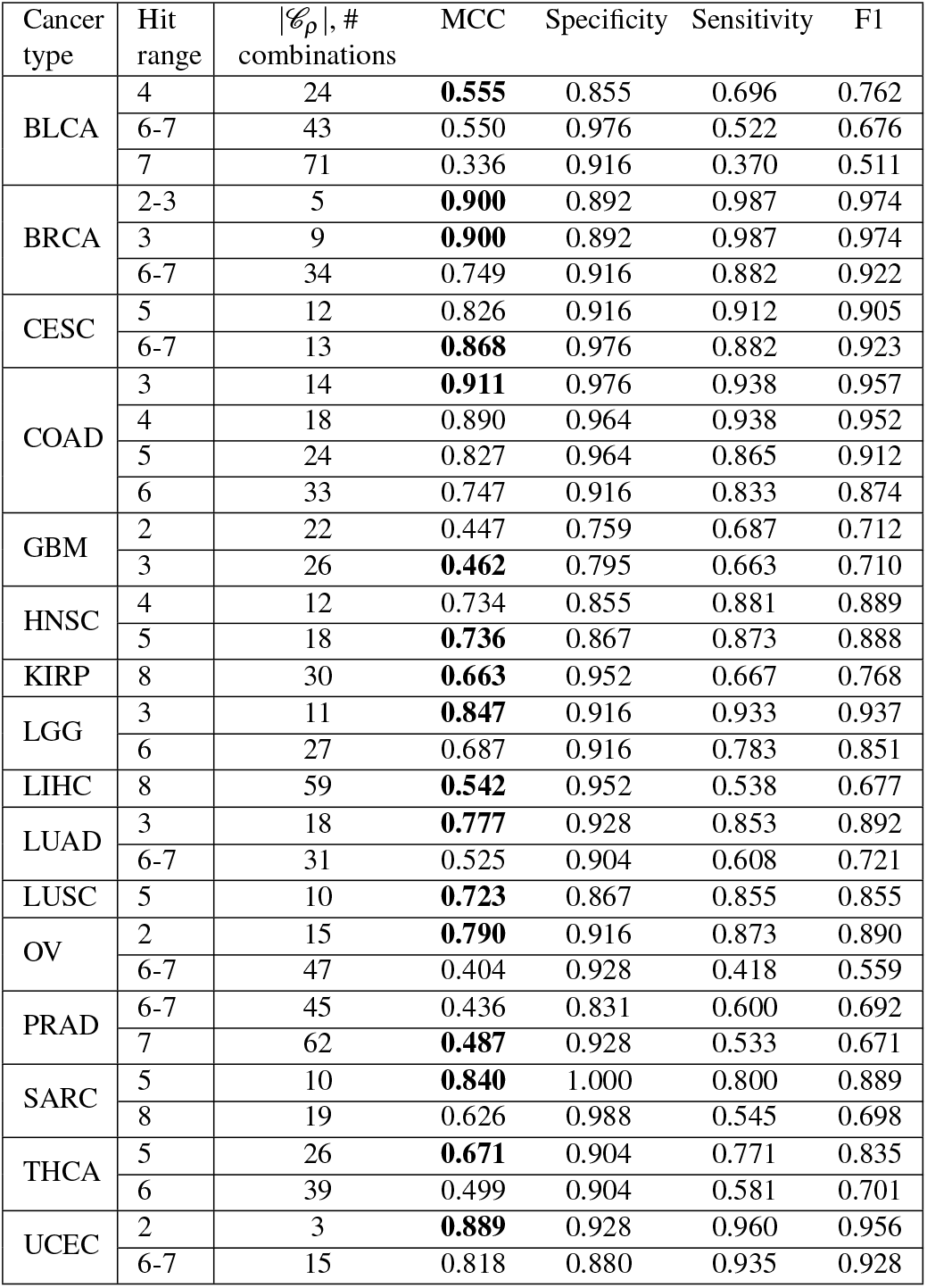
Classification performance of identified combinations on 𝒮_test_. Best MCC values per cancer type are in bold.

| Cancer type | Hit range | $\vert \mathcal{C}_\rho\vert , \#$<br>combinations | MCC | Specificity | Sensitivity | F1 |
| --- | --- | --- | --- | --- | --- | --- |
| BLCA | 4 | 24 | **0.555** | 0.855 | 0.696 | 0.762 |
|  | 6-7 | 43 | 0.550 | 0.976 | 0.522 | 0.676 |
|  | 7 | 71 | 0.336 | 0.916 | 0.370 | 0.511 |
| BRCA | 2-3 | 5 | **0.900** | 0.892 | 0.987 | 0.974 |
|  | 3 | 9 | **0.900** | 0.892 | 0.987 | 0.974 |
|  | 6-7 | 34 | 0.749 | 0.916 | 0.882 | 0.922 |
| CESC | 5 | 12 | 0.826 | 0.916 | 0.912 | 0.905 |
|  | 6-7 | 13 | **0.868** | 0.976 | 0.882 | 0.923 |
| COAD | 3 | 14 | **0.911** | 0.976 | 0.938 | 0.957 |
|  | 4 | 18 | 0.890 | 0.964 | 0.938 | 0.952 |
|  | 5 | 24 | 0.827 | 0.964 | 0.865 | 0.912 |
|  | 6 | 33 | 0.747 | 0.916 | 0.833 | 0.874 |
| GBM | 2 | 22 | 0.447 | 0.759 | 0.687 | 0.712 |
|  | 3 | 26 | **0.462** | 0.795 | 0.663 | 0.710 |
| HNSC | 4 | 12 | 0.734 | 0.855 | 0.881 | 0.889 |
|  | 5 | 18 | **0.736** | 0.867 | 0.873 | 0.888 |
| KIRP | 8 | 30 | **0.663** | 0.952 | 0.667 | 0.768 |
| LGG | 3 | 11 | **0.847** | 0.916 | 0.933 | 0.937 |
|  | 6 | 27 | 0.687 | 0.916 | 0.783 | 0.851 |
| LIHC | 8 | 59 | **0.542** | 0.952 | 0.538 | 0.677 |
| LUAD | 3 | 18 | **0.777** | 0.928 | 0.853 | 0.892 |
|  | 6-7 | 31 | 0.525 | 0.904 | 0.608 | 0.721 |
| LUSC | 5 | 10 | **0.723** | 0.867 | 0.855 | 0.855 |
| OV | 2 | 15 | **0.790** | 0.916 | 0.873 | 0.890 |
|  | 6-7 | 47 | 0.404 | 0.928 | 0.418 | 0.559 |
| PRAD | 6-7 | 45 | 0.436 | 0.831 | 0.600 | 0.692 |
|  | 7 | 62 | **0.487** | 0.928 | 0.533 | 0.671 |
| SARC | 5 | 10 | **0.840** | 1.000 | 0.800 | 0.889 |
|  | 8 | 19 | 0.626 | 0.988 | 0.545 | 0.698 |
| THCA | 5 | 26 | **0.671** | 0.904 | 0.771 | 0.835 |
|  | 6 | 39 | 0.499 | 0.904 | 0.581 | 0.701 |
| UCEC | 2 | 3 | **0.889** | 0.928 | 0.960 | 0.956 |
|  | 6-7 | 15 | 0.818 | 0.880 | 0.935 | 0.928 |

**Figure 2.**
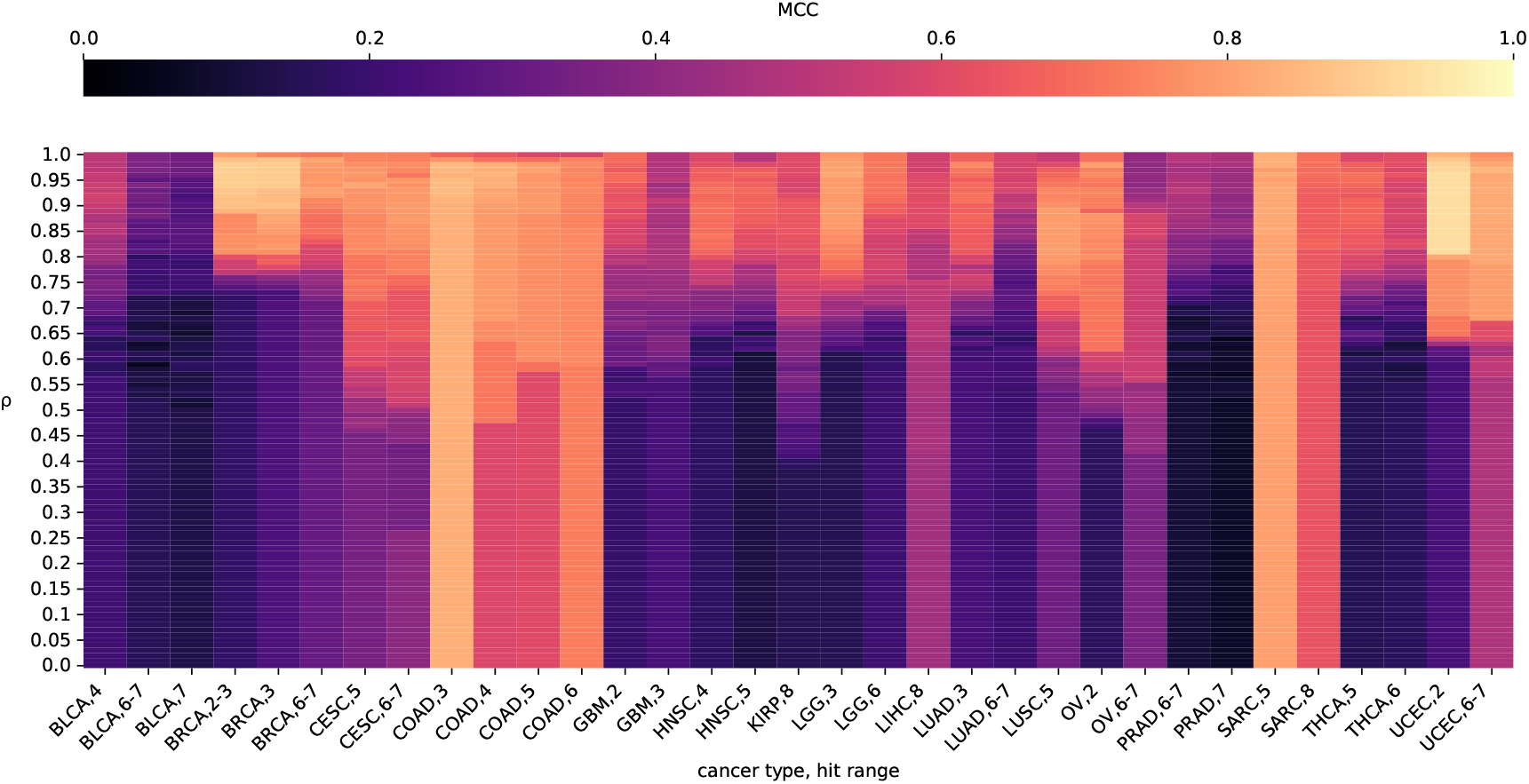
MCC fold-average of the MCC in cross-validation runs for the full range of *ρ* values. The range of MCC is truncated to [0, 1] due to the absence of negative values in our results.

### 3.3 Biological relevance

There are 316 genes from Cosmic gene database among the participating genes in the combinations identified by the full-dataset runs of BiGPICC. The number of Cosmic genes identified for different cancer types and hit ranges varies (Figure 3) but is generally proportional to the total count of identified genes in an experiment. For example, the number of Cosmic genes among the 7-hit combinations for BLCA, the worst performing classifier in our experiments, is 25 out of 303. At the same time, the “BRCA, 2-3” run has no Cosmic genes among the identified 10, despite delivering one of the highest classification performances. Because the Cosmic database contains the genes that are associated with cancer individually and BiGPICC searches for gene combinations, the dissonance between the Cosmic gene counts and classification performance can be partially attributed to the emergent carcinogenic properties of combined gene mutations. In addition, unlike many other studies BiGPICC does not set a minimum requirement for the number of explained tumor samples per combination which allows uncovering rare carcinogenic combinations. The potential presence of passenger genes in the combinations identified by BiGPICC (see Section 5) would also lower the respective proportion of Cosmic genes.

**Figure 3.**
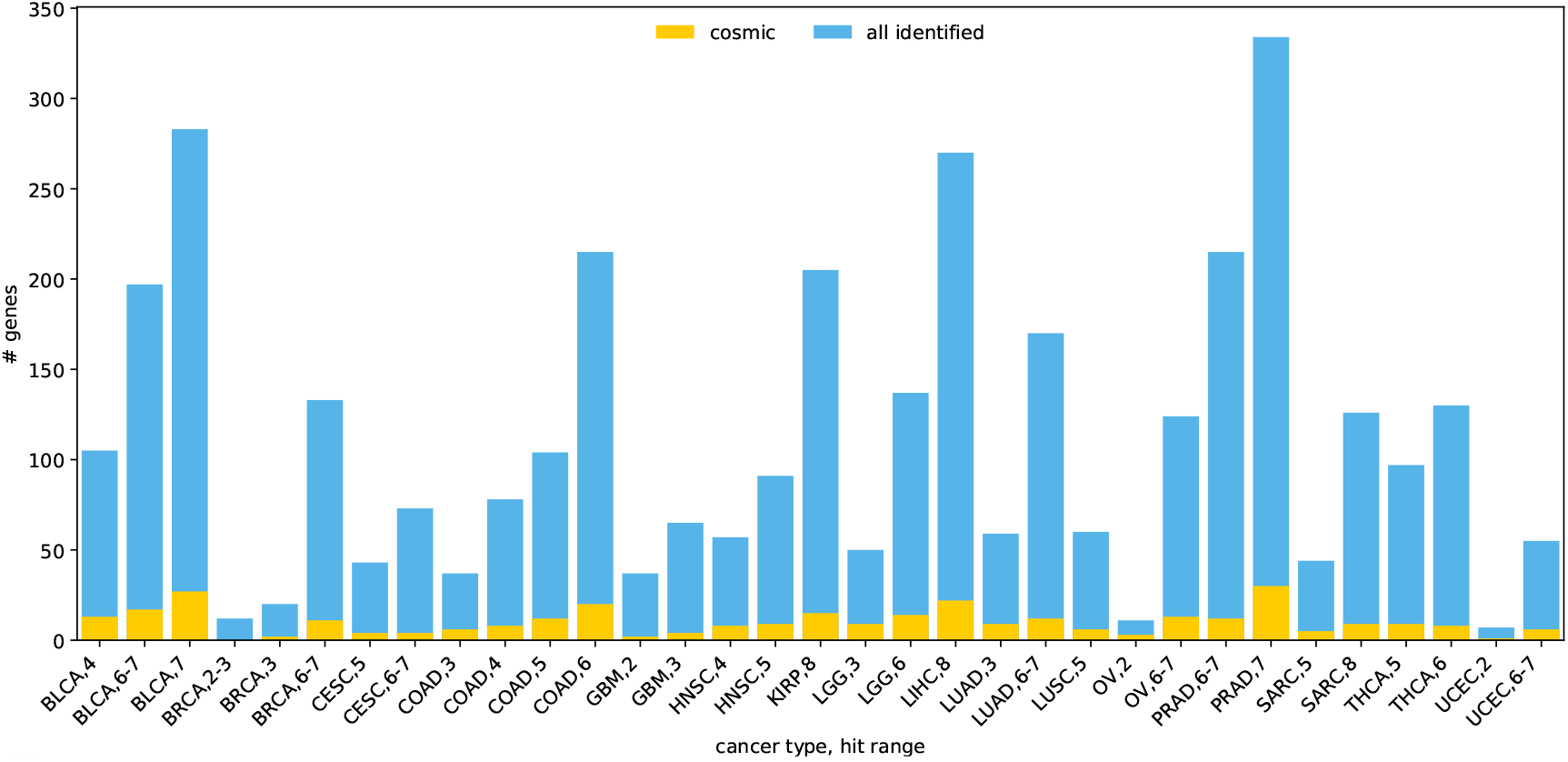
The proportion of Cosmic genes in all the carcinogenic genes identified by BiGPICC in each experiment.

### 3.4 Synthetic validation of the driver-passenger distinction

Trivially, passenger mutations that always co-occur with the driver mutations cannot be discerned from the latter using mutation data alone. To assess BiGPICC’s ability to tell apart passenger mutations with an arbitrarily high but non-100% chance of developing in the presence of driver mutations, we run an experiment on synthesized tumor-sample mutation data. We simulate mutations of 20,000 “genes” in 1,000 “tumor samples” with the baseline probability of 4.1% per sample — the average mutation rate of a gene in tumor samples from the 16 TCGA datasets. For 3 ≤ *h* ≤ 8, we choose *h* genes out of the 20,000 to represent the driving combination of carcinogenesis, and set them as mutated in 100% of the samples. Lastly, we designate another 1, *h*, 10*h*, or 100*h* genes as passenger and introduce their mutations with 99% probability in every sample.

Note that extending the data to include normal samples would not introduce new gene combinations and can only lower the chances of a particular combination to pass the tumor ratio-based filtration and thus to appear in the minimum cover. In particular, it means that combinations containing passenger mutations are most likely to appear in the output of BiGPICC when it is run on tumor-only data.

We apply the pipeline to the synthetic tumor-only dataset as defined above, running the community detection step (Procedure 1) *p* = 100 times in each experiment and setting the tumor ratio threshold to *ρ* = 1. Remarkably, in all 24 experiments (6 values of *h* paired with 4 passenger gene counts for each) the resulting minimum cover is comprised by a sole combination which is the correct carcinogenic one. At the same time, when the number of tumor samples is reduced to 500, the algorithm fails to identify the correct combination as the only driver of carcinogenesis in the experiments with 10*h* and 100*h* passenger genes, due to some passenger mutations appearing in every tumor sample. This demonstrates that our algorithm is capable of the driver-passenger distinction subject to availability of sufficient amounts of data ensuring the existence of unaffected tumor samples for every passenger mutation.

### 3.5 Comparison against exhaustive search results

The authors of Al Hajri et al.^18^ identified carcinogenic combinations of length 2, 3, and 4 for various cancer types through an exhaustive search (ES) of all possible gene combinations of given length. Similarly to the minimum set cover step of BiGPICC, they iteratively add carcinogenic gene combinations until their resulting set explains all tumor samples in the data. Each step of ES approach selects the combination *C*_𝒢_ maximizing *αn*_TP_ + *n*_TN_, where *n*_TP_ is the number of currently unexplained tumor samples in which *C*_𝒢_ is jointly mutated and *n*_TN_ is the number of normal samples in which it is not. The parameter *α* represents the importance of correctly identifying normal samples relative to the analogous importance for tumor samples, and was fixed to *α* = 0.1 in all ES experiments.

Figure 4 shows the comparison between the classification performance of carcinogenic combinations identified by BiGPICC for *h <* 5 and the metrics reported in Al Hajri et al. The two methods have comparable performance except for GBM cancer where BiGPICC performs significantly worse than ES. In seven out of ten cases, BiGPICC outperforms ES in either specificity or sensitivity without a significant compromise in the other metric, and in two cases — for both metrics. A possible explanation for the heuristics-based BiGPICC outperforming an exhaustive search method may be a suboptimal choice of constant *α* = 0.1 in the latter. In addition, the train-test splits used in our runs are not the same as in Al Hajri et al. If the amount of samples in the dataset is insufficient, a particular choice of the split may significantly impact the resulting performance, e.g. if the training samples 𝒮 \ 𝒮_test_ fail to capture all relevant gene interactions. In particular, our second attempt of the random train-test split of GBM data resulted in significantly higher MCC scores on 𝒮_test_ — 0.791 for GBM, 2 and 0.714 for GBM, 3, with the new specificity for GBM, 2 outperforming its ES counterpart.

**Figure 4.**
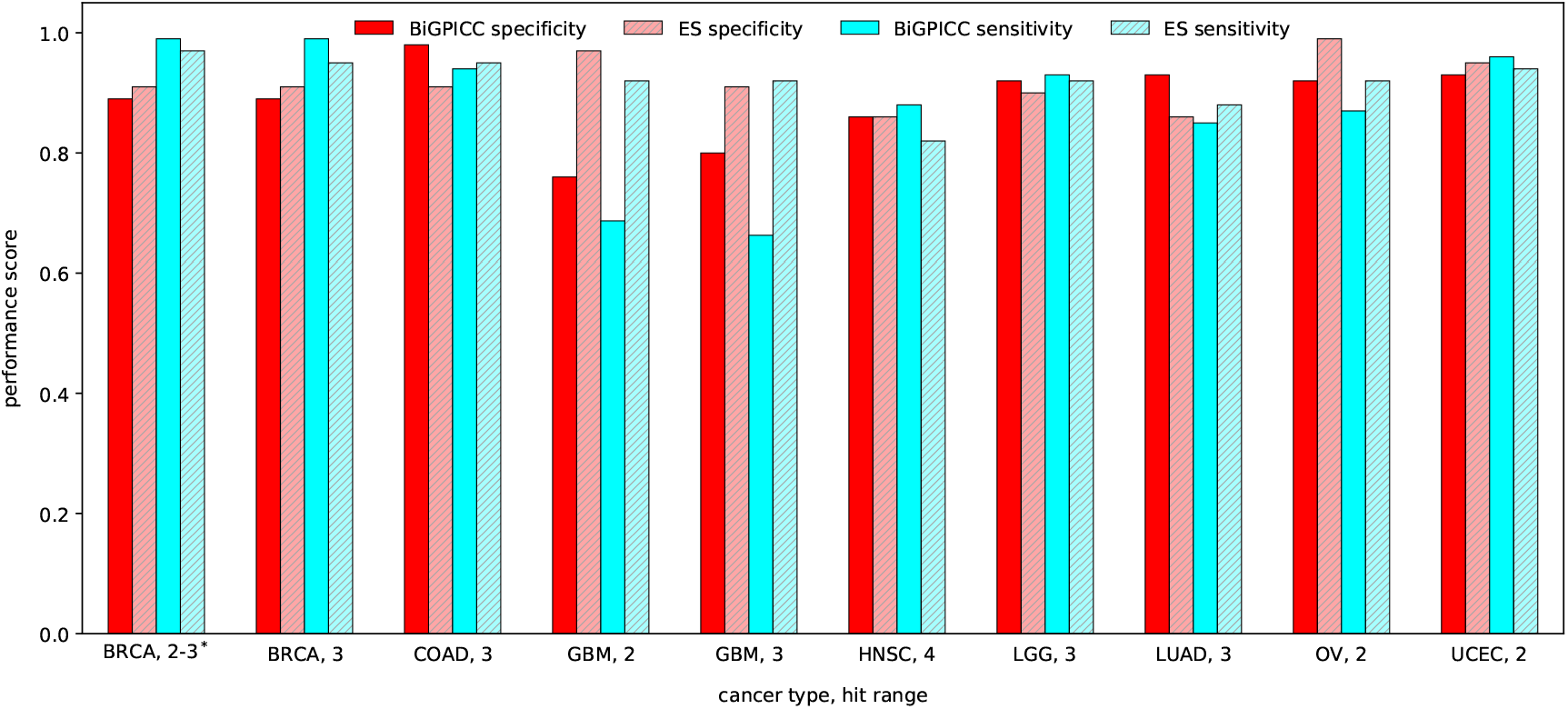
Comparison between classification performances of BiGPICC and Exhaustive Search (ES), rounded to 2 decimal places. ^*^BiGPICC runs for BRCA, 2-3 result in at most one 3-hit combination per fold and are compared against the ES results for BRCA, 2 as the best matching option.

### 3.6 Runtime performance

The 4-fold cross-validation BiGPICC runs on 75% of all samples for tuning the parameter *ρ* were conducted using 280 Summit nodes each and took on average 26 minutes across all experiments, ranging from 11 minutes (SARC, 8) to 1 hour 29 minutes (UCEC, 2). Running the pipeline on the full dataset (without the validation samples) to measure the resulting classification performance was done using 70 nodes and took 23 minutes on average, ranging from 8 minutes (SARC, 8) to 1 hour 33 minutes (UCEC, 6-7). Adding the two mean runtimes gives that identifying carcinogenic combinations on the full dataset from scratch takes BiGPICC on average 146 node-hours on Summit.

For comparison, the ES method from Dash et al.^17^ requires 124 Summit node-hours to merely identify 4-hit combinations for BRCA cancer type based on 75% of the available samples. Assuming ideal scaling of the method, the time required to identify 5-, 6-, and 7-hit combinations using all 4,600 nodes of Summit supercomputer would be 4.4 days, 39 years, and 107,000 years, respectively. In contrast, the BiGPICC runtimes are not noticeably affected by the combination size *h* and instead depend on the size and topology of the input graph determined by the cancer type. It follows that BiGPICC is orders of magnitude faster than ES for any *h* > 4.

While ES relies on GPUs, BiGPICC only uses CPU cores. Per Summit node, BiGPICC uses 42 CPU cores, while ES uses 6 × 5,120 = 30,720 GPU cores. Though CPU and GPU cores are not directly comparable, BiGPICC is using roughly 2% of each compute node and thus can be moved to a less expensive CPU-only cluster to achieve the same runtime.

## 4 Conclusions and discussion

We proposed a community detection-based approach for identifying carcinogenic combinations and demonstrated that its classification accuracy on the TCGA dataset is comparable to that for state-of-the-art. At the same time, BiGPICC enables discovery of (5+)-hit combinations intractable for exhaustive search methods even on most modern supercomputers.

In all cancer types considered, biological relevance of the identified combinations, measured as classification performance, tends to drop as the number of hits increases. A similar trend in the exhaustive search results for *h* ≤ 4 from Al Hajri et al.^18^ suggests that, aside from biological reasons, the issue may lie with the so-called curse of dimensionality in both the machine learning and combinatorial contexts. An increase in the number of hits means fewer samples with joint mutation of a gene combination exist in the data while the search space of possible combinations grows exponentially. Therefore, the number of samples required for finding carcinogenic combinations grows with the number of hits, while our runs use the same dataset for every multi-hit range.

The classification performance of BiGPICC exhibits a significantly higher variability in sensitivity (the percentage of correctly classified tumor samples) than in specificity (the analogous percentage for normal samples). The sensitivity and specificity scores across the experiments are distributed as 0.763 ± 0.175 (mean ± SD) and 0.911 ± 0.054, respectively. We attribute this trend to the implicit control exhibited over the specificity score by the parameter *ρ*, whose learned value is consistently high in the experiments (0.938 ± 0.051). Let *n*_TP_, *n*_FN_, *n*_TN_, and *n*_FP_ denote respectively the number of correctly predicted tumor samples, incorrectly predicted tumor samples, correctly predicted normal samples, and incorrectly predicted normal samples in the training dataset with the total number of samples *n* = *n*_TP_ + *n*_FN_ + *n*_TN_ + *n*_FP_. Parameter *ρ* ensures that the combinations selected for the classifier have sufficient tumor ratio, which translates into controlling its precision 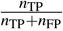 Assuming for simplicity no overlap between the samples in which the selected combinations are jointly mutated, the relationship is given by 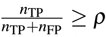 (in general, the right hand side can be both larger or smaller depending on the tumor ratio in the overlapping samples). It follows that the number of false positives is limited by the number of tumor samples in the dataset, 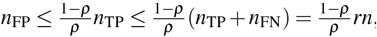, where 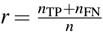 is the ratio of tumor samples in the data. Because *n*_FP_ is the only factor negatively affecting the specificity 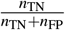, the latter is bounded from below by 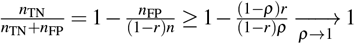. In particular, the average value of 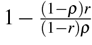 across the experiments is 0.865 (if *r* is calculated using the full datasets; it is expected to be the same along the training-validation split). Importantly, the pipeline does not exhibit control over *n*_FN_, the adverse factor for the sensitivity 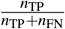, which is comprised by the tumor samples left unexplained by “sufficiently carcingonenic” combinations.

Unlike most other approaches, BiGPICC does not require manually balancing the importance of tumor and normal samples in the data. Instead, its hyperparameter *ρ* which implicitly takes on this role is learned from the data using cross-validation. This renders our approach parameter-free, alleviating the need to manually tune it on a case-by-case basis.

## 5 Limitations and future work

BiGPICC does not offer a dedicated mechanism to differentiate between the driver mutations causing carcinogenesis and the passenger mutations that do not contribute to cancer formation. Instead, background mutation rate typically used for such differentiation (e.g. in ^20^) is implicitly accounted for at the stage of identifying candidate combinations, as more frequently co-mutating genes form denser communities (see the discussion of unrelated genes at the end of 2.1). However, if the available samples are insufficient for such a differentiation, some of the genes in the identified combinations can lack biological relevance (due to mutating in tumor samples by chance), thus compromising specificity, but not sensitivity of the method. In particular, each of our datasets contains between 41 and 67 genes with mutations in over 50% of the samples. This can lead to many candidate combinations with near-identical explanatory power, which can be behind the observed phenomenon of the similar classification performance exhibited by significantly different sets of carcinogenic combinations. Increasing the number of samples in the data would mitigate the issue of driver-passenger distinction by amplifying the signal of carcinogenic pathways against the backdrop of noise from chance mutations. Under the assumption that highly mutable genes are unlikely to drive carcinogenesis, another approach would be to remove genes with mutation frequency above some domain-informed threshold from the analysis — we avoided doing so in this study in order to demonstrate the algorithm’s performance in the absence of tuning to a particular set of data.

Another limitation of the method is that it is run on the gene-patient data encoding all possible mutations of a gene as a binary variable, thus ignoring the variability in mutations of individual genes. However, the BiGPICC pipeline could similarly be applied to mutation-level data for the identified carcinogenic gene combinations. Given that the identified genes typically constitute less than 1% of the original gene pool, we expect mutation-level data to be of similar size and tractability to the datasets used in this study. Incorporating additional omics data, such as transcriptomics, can enhance the comprehensiveness and relevance of identifying mutations within a gene by providing functional insights into their impact. Applying our approach to the resulting mutation-level data would be an important next step towards efficient diagnostics and drug discovery.

Assuming that the sufficient amount of data is available, an additional use of our pipeline is testing out the competing ranges of multi-hit theories by comparing the performance of learned carcinogenic combinations of the corresponding size. The ICGC database^36^ contains roughly twice as many samples as our datasets for individual cancer types, and may be a prospective dataset to try this approach on.

## Declarations

## Ethics approval and consent to participate

Not applicable.

## Consent for publication

Not applicable.

## Availability of data and material

Our code and the datasets for two cancer types are available at https://code.ornl.gov/vo0/bigpicc. The datasets for other cancer types are available from Sajal Dash upon request.

## Author contributions

R.A. preprocessed the data, S.D. conceived the idea, V.O. designed and implemented the algorithm, V.O. and S.D. ran the experiments and analyzed the results. All authors reviewed the manuscript.

## Acknowledgements

This work was supported by the resources of the Oak Ridge Leadership Computing Facility, located in the National Center for Computational Sciences at ORNL, which is managed by UT Battelle, LLC for the U.S. DOE (under the contract No. DE-AC05-00OR22725).

